# Systematic Replication Enables Normalization of High-throughput Imaging Assays

**DOI:** 10.1101/2022.04.26.489617

**Authors:** Gregory J. Hunt, Mark A. Dane, James E. Korkola, Laura M. Heiser, Johann A. Gagnon-Bartsch

## Abstract

**Motivation:** High-throughput fluorescent microscopy is a popular class of techniques for studying tissues and cells through automated imaging and feature extraction of hundreds to thousands of samples. Like other high-throughput assays, these approaches can suffer from unwanted noise and technical artifacts that obscure the biological signal. In this work we consider how an experimental design incorporating multiple levels of replication enables removal of technical artifacts from such image-based platforms.

**Results:** We develop a general approach to remove technical artifacts from high-throughput image data that leverages an experimental design with multiple levels of replication. To illustrate the methods we consider microenvironment microarrays (MEMAs), a high-throughput platform designed to study cellular responses to microenvironmental perturbations. In application on MEMAs, our approach removes unwanted spatial artifacts and thereby enhances the biological signal. This approach has broad applicability to diverse biological assays.

**Availability:** Raw data is on synapse (syn2862345), analysis code is on github (gjhunt/mema norm), a Docker image is available on dockerhub (gjhunt/memanorm). online.

## 1 Introduction

Experimental designs incorporating replication and randomization can significantly empower the analysis of large scale datasets and facilitate their biological interpretation. Replication, in particular, has a multitude of well-known benefits. Replication can be used to detect and understand sources of error, for example by comparing replicates in an exploratory analysis. It can also help quantify random variation, e.g., through calculation of standard deviations. Furthermore, replication can also help mitigate errors, for example by averaging over replicates. Indeed, replication can pay particular dividends in the analysis of high-dimensional biological data, where it can be used to identify and remove unwanted systematic effects. In this paper we demonstrate these principles by showing how careful experimental design can enable normalization of imaging assays. Our approach takes advantage of novel opportunities for normalization by exploiting multiple levels of replication within the experimental design, as well as by sharing knowledge across the hundreds of biological features that are extracted from each individual image.

The approach we develop focuses on imaging assays that make use of fluorescent microscopy. Fluorescent microscopy is a class of optical techniques to study the cellular and phenotypic responses of tissues and cells (Heilemann, 2012). These techniques image cells that have been labelled with fluorescent molecules designed to highlight cellular and extracellular structures of interest. The wealth of high-quality image data that can be created with such techniques has made it popular across the life sciences (Sood *et al*., 2020; Hamilton, 2009; Ishizawa et al., 2009; Huwyler and Pardridge, 1998). For example, immunohistochemistry and immunocytochemistry biomarkers can be used to highlight biologically-meaningful proteins that enable assessment of cell morphology, pathway activity and differentiation state (Drew and Shieh, 2015). More recently, highly-multiplexed techniques, including multiplexed immunofluorescence (MxIF), cyclic immunofluorescence (CycIF) and multiplexed immunohistochemical consecutive staining on single slide (MICSSS) have enabled assessment of tens of markers through a process of repeated of staining, imaging, and dye inactivation (Tan *et al*., 2020; Lin et al., 2016; Akturk et al., 2020; Bray et al., 2016). These approaches allow a wide array of structures to be simultaneously visualized in a single sample. Additionally, high-throughput experiments that use automated staining and imaging allow processing of hundreds to thousands of samples over the course of an experiment. The large volume of data produced by these techniques requires quantitative analysis with automated or semi-automated image analysis platforms such as CellProfiler, ImageJ, QuPath, or Ilastik (Carpenter *et al*., 2006; Collins, 2007; Bankhead *et al*., 2017; Sommer et al., 2011). Such platforms enable extracting hundreds of quantitative features from the images at both the cell and tissue level.

The present work focuses on microenvironment microarrays (MEMAs), an image-based cell-profiling approach that combines high-throughput fluorescent microscopy and spotted arrays (Lin *et al*., 2012; Smith *et al*., 2019; Watson *et al*., 2018). In brief, insoluble proteins are printed in a grid-like manner on a tissue culture plate to form pads upon which cells can be grown. Soluble factors are added to the culture medium, which together with the printed proteins yield thousands of unique combinatorial microenvironment perturbagens. The cells are subsequently fixed, stained, imaged, and analyzed to assess the impact of microenvironmental factors on cellular morphology and phenotype.

Such imaging data can suffer from unwanted technical effects, which challenges their biological interpretation. For example, unwanted effects include variations in staining intensity, background autofluorescence, and image alignment or segmentation issues can induce unwanted systematic artifacts in the data. Consequently, a small number of data normalization approaches have been developed to address some of these sources of unwanted effects for particular imaging pipelines (Chang *et al*., 2020; Ahmed Raza *et al*., 2016; Harris *et al*., 2021). MEMAs are a relatively new assay and to date there remains no general approach to remove unwanted effects from MEMA image data. However, their array-based structure presents an opportunity to enable data normalization through experimental design. In this work we present a general approach to normalization that makes use of replication at multiple levels of the design. We show that our approach can help identify and remove unwanted effects without the need of external controls or known sources of the effects. Furthermore, we show that this normalization improves detection of the desired biological signals. While performance is demonstrated on the MEMA assay, the design considerations and normalization technique have broad applicability to many forms of high-throughput biological data.

## 2 Design and Methods

A successful normalization method must separate wanted biological signal from unwanted systematic effects. Our approach is to design the experiment from the start to help make these separable. In particular, we make use of a design that incorporates both (1) replication at multiple levels and (2) randomization.

Replicated measurements can help enable identification of unwanted effects. This is possible as, absent any unwanted effects, measurements across replicates should not differ. For example, if we run the same experiment twice, absent any unwanted effects, we expect the measurements from the two experiments to be the same. In a more realistic case, we may have unwanted effects and consequently the replicated measurements will not be equal. However, as any observed difference is necessarily driven by the unwanted effects we can use these differences to partially identify them.

For this strategy to succeed, there must be sufficient replication in the experimental design to capture all of the unwanted effects we wish to remove. Therefore, we developed an experimental design that builds in replication at several levels. In particular, our MEMA design includes multiple replicates at two levels: (1)we have several replicates of each of the extra-cellular matrix proteins (ECMps) in each well, and (2) we have multiple replicates of each soluble ligand. These two levels of replication will each provide distinct information about the undesired effects. Additionally our design randomizes the spatial layout of the ECMps in the wells to ensure that the unwanted effects vary over the replicates. In the subsequent section we introduce the specific experimental design of the data we will analyze. After describing the design and data, Section 2.2 will present a method that leverages the design to find and remove unwanted effects. In doing so, our approach underlines the power of incorporating a robust experimental design into the planning of high-throughput imaging experiments.

### 2.1 Experimental Design

A microenvironment microarray (MEMA) is a spot-based assay used to study perturbations to the physical and chemical surroundings of cells. Figure 1 (A) displays the MEMA data-generation pipeline. Here, we show a MEMA plate which consists of a plastic tissue culture plate divided into eight wells. First, ECMps are printed into each of these wells in a grid of 20 by 35 spots. Typically 45 unique ECMps are printed in 15 replicates. After printing the ECMps, cells are added to each spot and a buffer solution is added to the wells. In addition to the buffer solution, soluble ligands are added to a selection of wells. A different ligand can be added to each well. After plating, the cells are grown for 72 hours. Subsequently fluorescently labelled antibodies or stains are applied and the cells are imaged. These images are then segmented and features are extracted. The extracted features capture a wide range of morphological aspects such as cell size (number of pixels), perimeter, eccentricity (measuring cell elongation), and many others. The extracted features also capture the intensity of the stains in the nucleus and cytoplasm of the cells.

**Figure 1:**
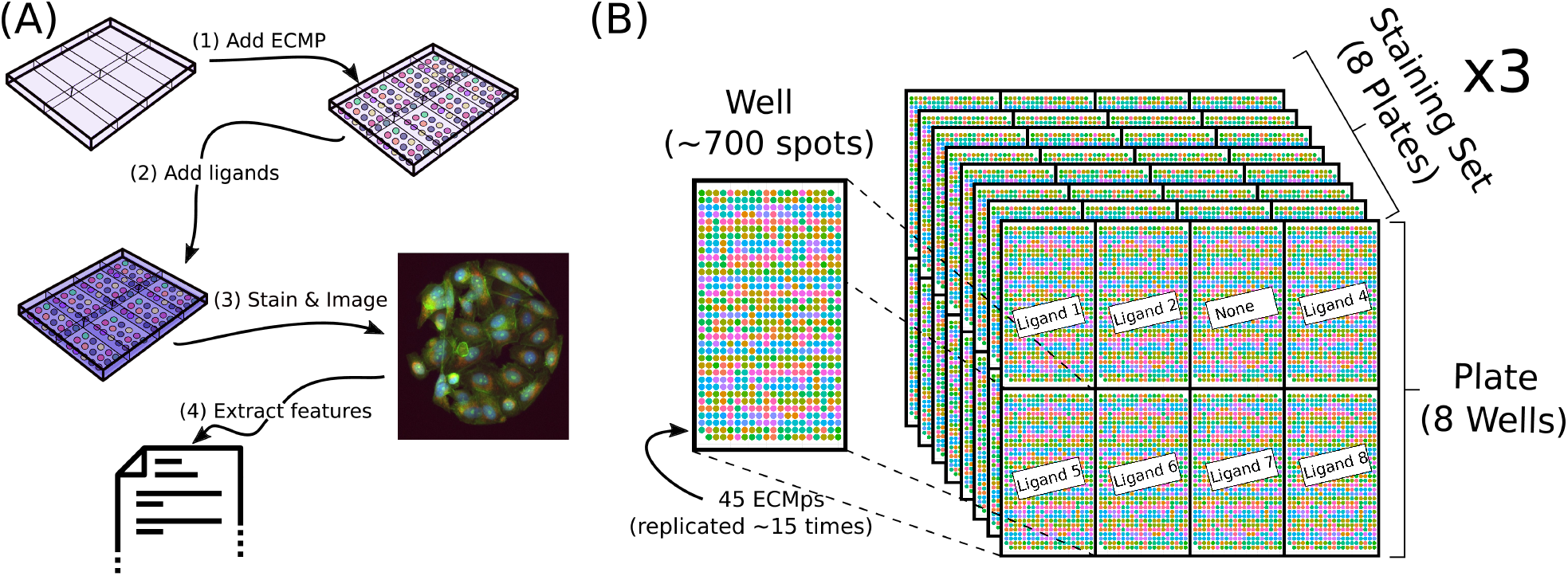
(A) Our data-generation pipeline for MEMAs. The MEMA substrate is sub-divided into eight wells, each containing a grid of approximately 700 spots comprised of a panel of 45 insoluble extracellular matrix proteins (ECMps). Once printed, cells are added to the well where they bind to the ECMp spots. Subsequently, soluble ligands are added to the buffer solution of each well. After growing, the cells they are stained and imaged with fluorescence microscopy. Finally, automated image analysis software is used to extract features from the images. (B) Experimental design. Each well contains approximately 700 spots with 45 ECMps replicated about 15 times each. This well design is replicated eight times across the wells in each plate with different ligands per well. Eight plates with different ligands applied to the wells of each plate constitute a staining set. The same set of three stains are applied to each of the plates in a staining set. There are three staining sets with different stains and hence different features extracted across the staining sets.

An important aspect of MEMAs is that ECMps vary from spot-to-spot and the ligands vary from well-to-well. Consequently, a MEMA allows one to investigate the effects of several thousand combinations of ECMps and ligands. A key to our approach is ensuring enough replication and randomization across these combinations to identify unwanted effects. We build in replication at two levels: (1) ECMps are replicated across spots, and (2) ligands are replicated across plates; we build in randomization at one level: the physical distribution of ECMps across spots follows a randomized design.

Figure 1 (B) displays a schematic of the experimental design. The left side displays the wells with color of the approx. 700 spots indicating one of 45 ECMps each replicated approximately 15 times. The ECMp pattern was designed with an initial one-time randomization and is the same across all wells. Several spots have no ECMp to guide physical orientation. The right side of Figure 1 (B) displays each plate consisting of two rows of four wells. Each well gets a different ligand treatment (including one with no ligand). Our data contains 24 plates processed in three batches of eight. These three batches are identically designed other than the stains applied. Correspondingly, these batches are called “staining sets” because the plates in each set have the same stains applied but plates across different sets may have different stains. All of the sets are stained with DAPI to visualize the nucleus and then three more stains are applied to each set. These three other stains vary from one set to another. The common DAPI stain localizes to the cell nucleus and allows us to segment the cells and extract a common set of morphological image features such as nuclear area and shape. Since the other stains are not applied to all staining sets the features derived from these stains are only captured for wells/plates in these corresponding sets.

Nonetheless, we will show that the replication of DAPI across all wells allows us to learn a normalization transformation on the DAPI-derived features and apply this normalization to features derived from other stains.

### 2.2 Normalization Method

As different features may be on different measurement scales, we pre-process our data using a three-step transformation which (1) Gaussianizes the data with a Box-Cox procedure, (2) robustly *z*-scores the features, and then (3) removes any extreme outliers (Hunt *et al*., 2020). This brings the disparate features to comparable scales, which is a critical step for our approach.

We work with our data in a well-by-spot matrix format. Let *n* denote the total number of wells across our experiment and *p* denote the number of spots per well. In our case *n* = 192 (8 wells *×* 8 plates *×* 3 staining sets) and *p* = 667. Each of the wells is associated with one of *ℓ* = 57 ligand conditions (either no ligand or one of the 56 added ligands). Each of the spots has been treated with one of *e* = 45 ECMps. After staining and imaging, image features are extracted using Cell Profiler. Precisely which features are extracted depends on the staining set with the exception of the DAPI-derived features which are extracted in all cases. In all, for the *i*^*th*^ feature we have an *n × p* matrix *Y* ^*i*^ encoding the values of the feature.

Note that the non-DAPI *Y* ^*i*^ will have systematically missing rows since some of the wells were not part of a staining set that allowed identification of the particular feature. For example, one-third of the wells (rows) for features derived from the Cell Mask stain will be entirely missing since only two out of the three staining sets had this stain. Without loss of generality, assume that the first *c* features 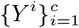 are the DAPI-derived features measured in all three staining sets and thus have no systematic missingness. We will call these the complete feature matrices.

The systematic missingness inherent to the data means that for some features the ECMp/ligand replicate structure among non-missing values is systematically incomplete. This is important because the replicate structure is what will allow us to separate the unwanted effects from the underlying biological signal. Later, we will discuss how we can leverage the complete features to remove unwanted effects from all features, even those with systematically missing values.

Without unwanted effects we assume that our data follows the model

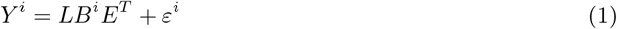

for a random *ε*^*i*^. Here, *L* encodes the ligand condition so that it is a *n ×. ℓ* indicator matrix of 0s and 1s where *L*_*jk*_ = 1 if well *j* is treated by the *k*^*th*^ ligand condition and zero otherwise. Similarly, *E* is the *p × e* ECMp indicator matrix so that *E*_*jk*_ = 1 if the *j*^*th*^ spot in the well is treated by ECMP *k*, and zero otherwise. Finally, let *B*^*i*^ be the corresponding. *ℓ × e* matrix that encodes the true biological signal of the ligands and ECMps for the *i*^*th*^ feature so that 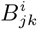 is the response of the *j*^*th*^ ligand and the *k*^*th*^ ECMp on feature *i*.

The model proposed by Equation 1 simply expands out the true ECMp-by-ligand signal encoded by *B*^*i*^ according to the experimental design encoded in *L* and *E*. While this proposed structure has similarities to a linear model, in our case the elements of *B*^*i*^ can vary arbitrarily and there is no assumed relationship or structure among the elements of *B*^*i*^. This makes the presumed model very general.

Unfortunately, our measurements captured in *Y* ^*i*^ likely suffer from systematic unwanted effects. Consequently, we augment our model so that

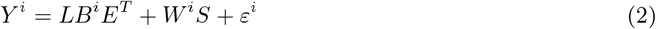

where *S* has the dimensions *d × p* and *W*^*i*^ is *n × d*. We can understand *S* as a matrix whose rows encode *d* unwanted effects over the *p* spots in a well. Correspondingly, each column of *W*^i^ encodes the extent to which the unwanted effects of *S* affect each of the *n* wells of *Y* ^*i*^. To remove the unwanted effects in our data we will form estimates *Ŵ* ^*i*^ and *Ŝ* and subtract *Ŵ* ^*i*^*Ŝ* from *Y* ^*i*^.

#### 2.2.1 Estimating *S*

For a matrix *A* let *RA* = *I − A*(*A*^*T*^ *A*)^*−*1^*A*^*T*^ be the residualizing operator that projects a vector *x* onto the orthogonal complement of the column space of *A*. That is, *R*_*A*_*x* is the residual from regressing *x* onto *A*. Notice that for our model

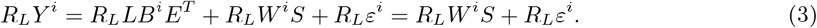

That is, if we residualize across the ligand replicates the *B*^*i*^ term disappears leaving only the unwanted effects and a random error term. Since this is true for all of the matrices, then consider residualizing the complete feature matrices 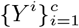 and stacking them, to form a *nc × p* matrix *Y*_*L*_ such that

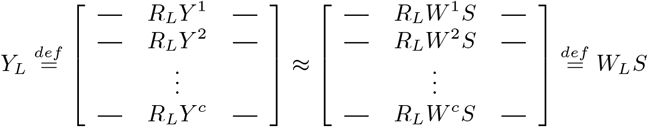

where *W*_*L*_ is defined so that the *i*^*th*^ row of *W*_*L*_ is *R*_*L*_*W*^*i*^. This suggests that we can estimate *S* with factor analysis on *Y*_*L*_. To do this we decompose *Y*_*L*_ with the SVD as *Y*_*L*_ = *W*_*L*_*S* = *U* S*V* ^*T*^ and estimate *S* as the first 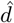 right-singular vectors of *Y*_*L*_ so that

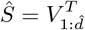

where *U*∑ correspondingly captures *W*_*L*_.

One can interpret our estimate of *S* as the eigenvectors of the average gram matrix of the *R*_*L*_*Y* ^*i*^. To see this recall that the right singular vectors *V* of *Y*_*L*_ are the eigenvectors of 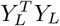. Now,

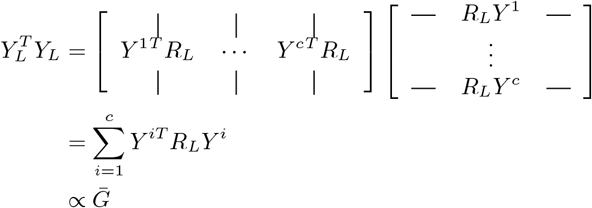

where we define *G*_*i*_ = *Y* ^*iT*^ *R*_*L*_*Y* ^*i*^ as the gram matrix of *R*_*L*_*Y* ^*i*^ and 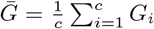 as the average. Thus we may alternatively think of *Ŝ* as formed from calculating 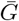, eigendecomposing it as

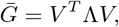

and taking the first 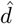 eigenvectors in *V* as *Ŝ*. This is how we will do it in practice. Because of missing values that may be present in the data our eigenvectors are computed using re-normalized completed gram matrices whose entries are computed using only the non-missing values (Hunt *et al*., 2020).

#### 2.2.2 Estimating *W*^*i*^

Now for any feature *Y* ^*i*^ consider residualizing to the ECMp replicate structure *E*. In this case,

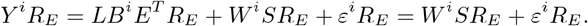

Define *S*_*E*_ = *SR*_*E*_ as the *d × p* matrix of *S* residualized to *E*. Notice that if *S*_*E*_ is full rank then 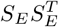 is invertible. Furthermore,

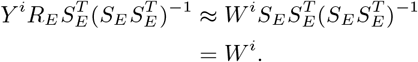

Thus for any *i*, even one for which the data has systematic missing values, we replace *S*_*E*_ with 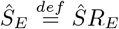 in the above equation and define our estimate of *W* as

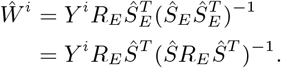

One can interpret *Ŵ* ^*i*^ as the regression coefficients corresponding to *Ŝ* if we were to regress *Y* ^*i*^ onto *Ŝ* and *E* simultaneously (Frisch and Waugh, 1933). This means that the efficiency of the estimate *Ŵ* ^*i*^ increases as the correlation between *E* and *Ŝ* decreases. In our design *E* follows a randomized design and so we expect this correlation to be low and thus *Ŵ* ^*i*^ to be efficient.

We summarize the full normalization approach in Algorithm 1.

##### Algorithm 1

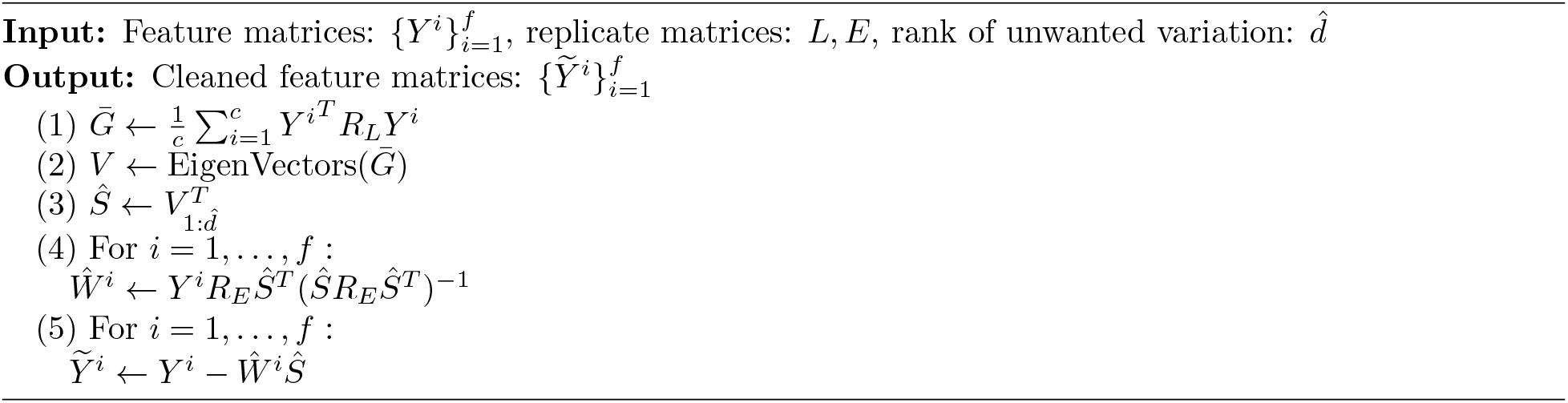

## 3 Results

In this section we assess the application of our approach to the MEMA data discussed in Section 2.1. We show that the method helps remove unwanted spatial effects present in the MEMA data and thereby enhances the recovery of the biological signal.

### 3.0.1 Common Spatial Patterns

One example of unwanted systematic effcts that can affect MEMAs and other image-based experiments is systematic spatial effects. For example, in the case of MEMAs, these unwanted spatial effects may be the result of imperfections in the plating of cells or ECMps or result from an unequal distribution of stain or ligands within the wells. Figure 2 displays several such spatial patterns. As the ECMp layout of the wells is randomized, we do not expect to see any such patterns.

**Figure 2:**
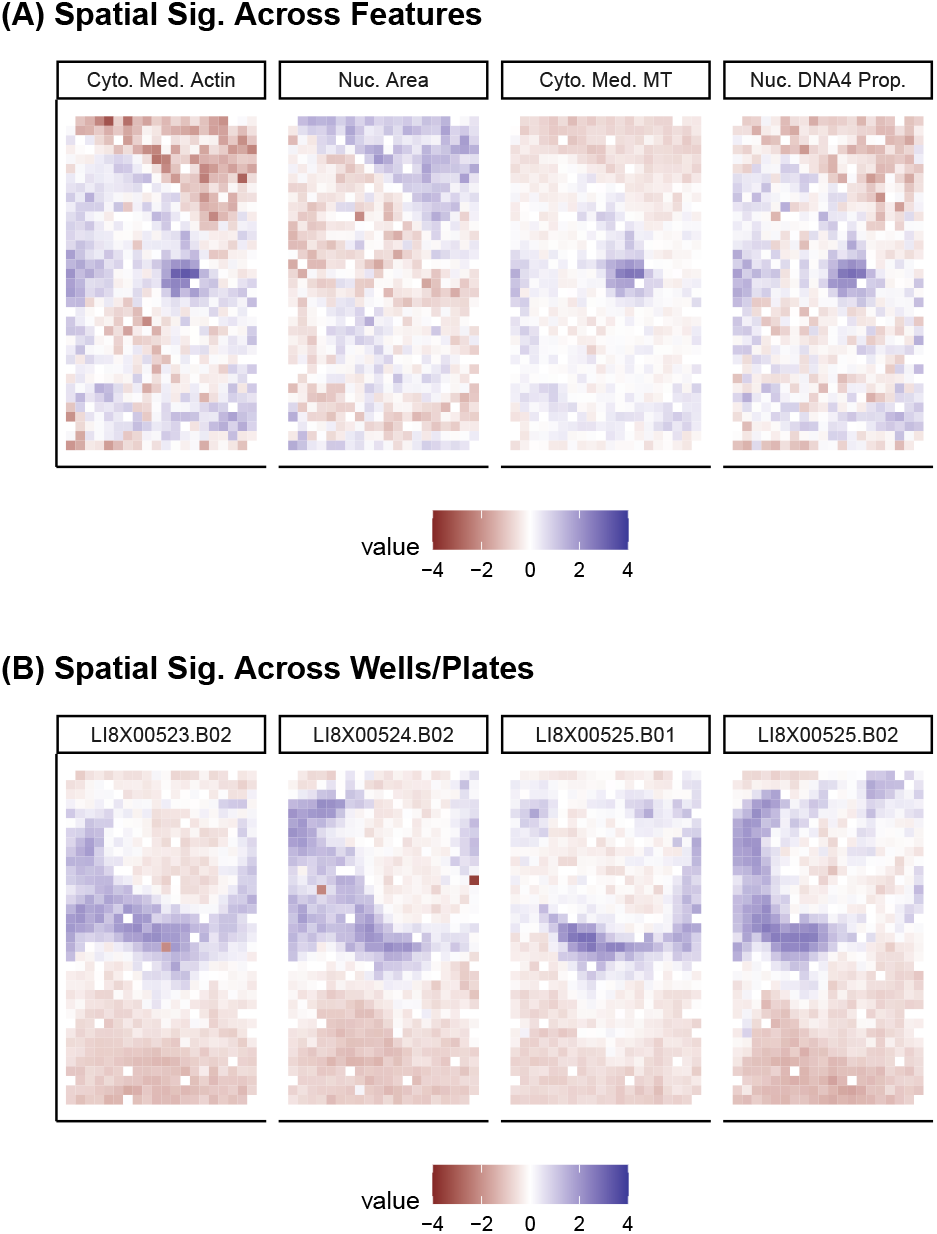
(A) Same spatial patterns across the features (1) median cytoplasmic Actin stain intensity, (2) nuclear area, (3) median cytoplasmic MitoTracker stain, and (4) nuclear DNA4 proportion, all in the same well. We can see a similar spatial patterns across these four features for this well. (B) Median cytoplasmic Actin stain intensity plotted for four different wells across three different plates. We can see a similar spatial pattern across these four wells even through they are on different plates.

The approach we outlined in Section 2 assumes that these unwanted spatial effects are shared across both the wells and features. This is modeled in Equation 2 by the low-dimensional *S* which is not indexed by *i* but shared among all features. The fact that the spatial effects are shared means we can estimate *S* using the features with a complete replication design and then remove these spatial effects from features with systematically incomplete replication. Note that this assumption is not as strong as it may first seem. Our approach does not require that the exact same unwanted effects show up in all features. Instead it makes a weaker assumption that there is some common space of spatial effects generally encompassing those encountered across the wells and features. The term *S* is precisely the basis for this space. Correspondingly, the model allows different features and wells to experience different unwanted effects through the the values of *W*^*i*^ which modulate the extent to which each of these spatial effects appear in any particular feature/well. Nonetheless, the ability to capture the spatial effects in a low-dimensional *S* does rely on the spatial effects broadly being shared across wells and features.

To show that these assumptions are reasonable for the present data, Figure 2 (A) displays an illustrative heat-map of four different image features over the spots of a chosen well. The features are: (1) median cytoplasmic actin stain intensity, (2) nuclear area, (3) median cytoplasmic MitoTracker stain, and (4) nuclear DNA4 proportion. In all four of these heat-maps we can broadly see a similar upper-right triangular pattern and also notice a dark blue artifact in the center of the wells, which indicates spatial effects in these image features. In Figure 2 (B) we display similar heat-maps as in subplot (A) but now focus on only one feature: the median cytoplasmic MitoTracker stain intensity. Here, we display four heat-maps corresponding to four different wells on three different physical plates. While the spatial pattern for these wells is different from subplot (A), we again see similar large-scale effects across the wells. In this case we notice a dark blue annular pattern. Notably, our approach does not assume any level of smoothness of the unwanted effects and can handle effects with a high degree of complexity as we see in these figures.

### 3.0.2 Spatial Effects Can Be Removed

To remove the unwanted spatial effects we use our procedure described in Section 2 and remove 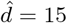 dimensions. In Figure 3 we display the basis for this space which comprises the columns of *Ŝ*. We can see from this figure that the unwanted effects we discover is largely spatial in nature for this particular experiment. This comports with our observations in Figure 2.

**Figure 3:**
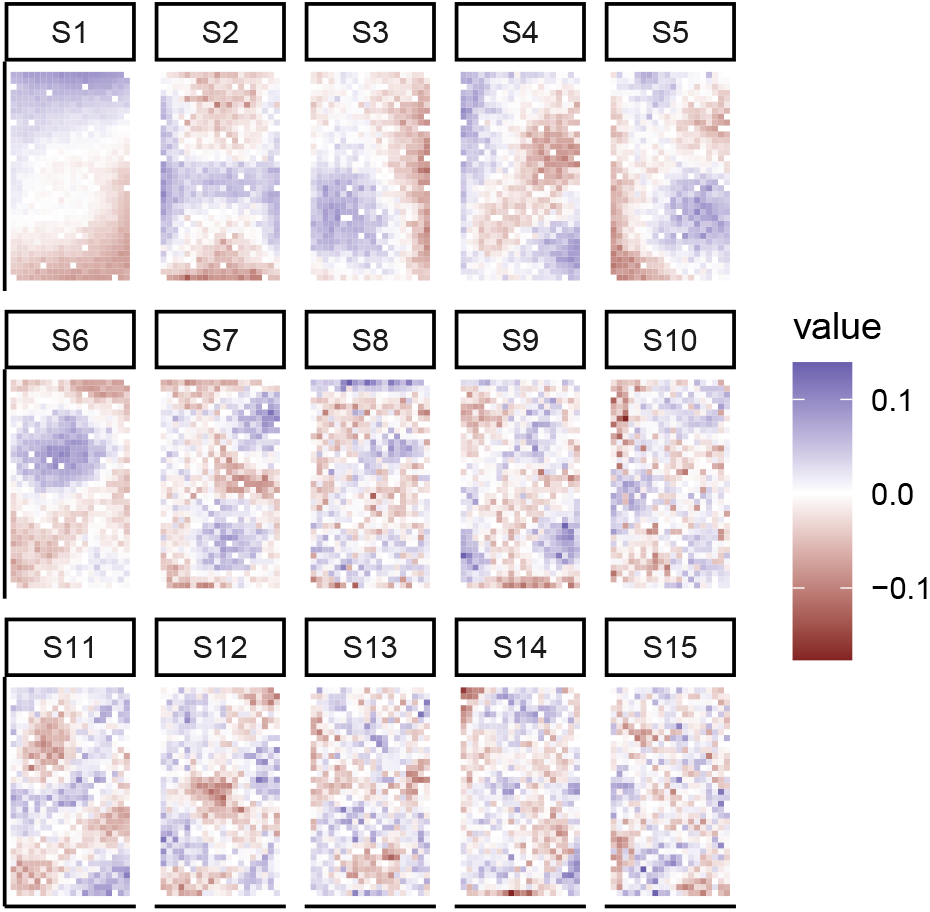
Unwanted signal extracted from the data using the replicate structure and our approach. This unwanted signal captures a low-dimensional representation of the spatial artifacts present in the MEMAs.

In Figure 4 (A) and (B) we replicate Figure 2 after removing the unwanted effects. We can see from these subplots that applying our procedure to these wells largely removes the spatial patterns. This is true even through the spatial patterns observed were very different in nature between subplots (A) and (B). This is possible through our estimates *Ŵ* ^*i*^ which determine the correct linear combination of the unwanted effects to remove. Note that most of the features in Figure 4 are not DAPI-derived and thus are not fully observed across the entire set of replications. Nonetheless, the spatial effects learned via the DAPI-derived features sufficiently capture those unwanted effects present in features derived from other stains that we are still able to remove them.

**Figure 4:**
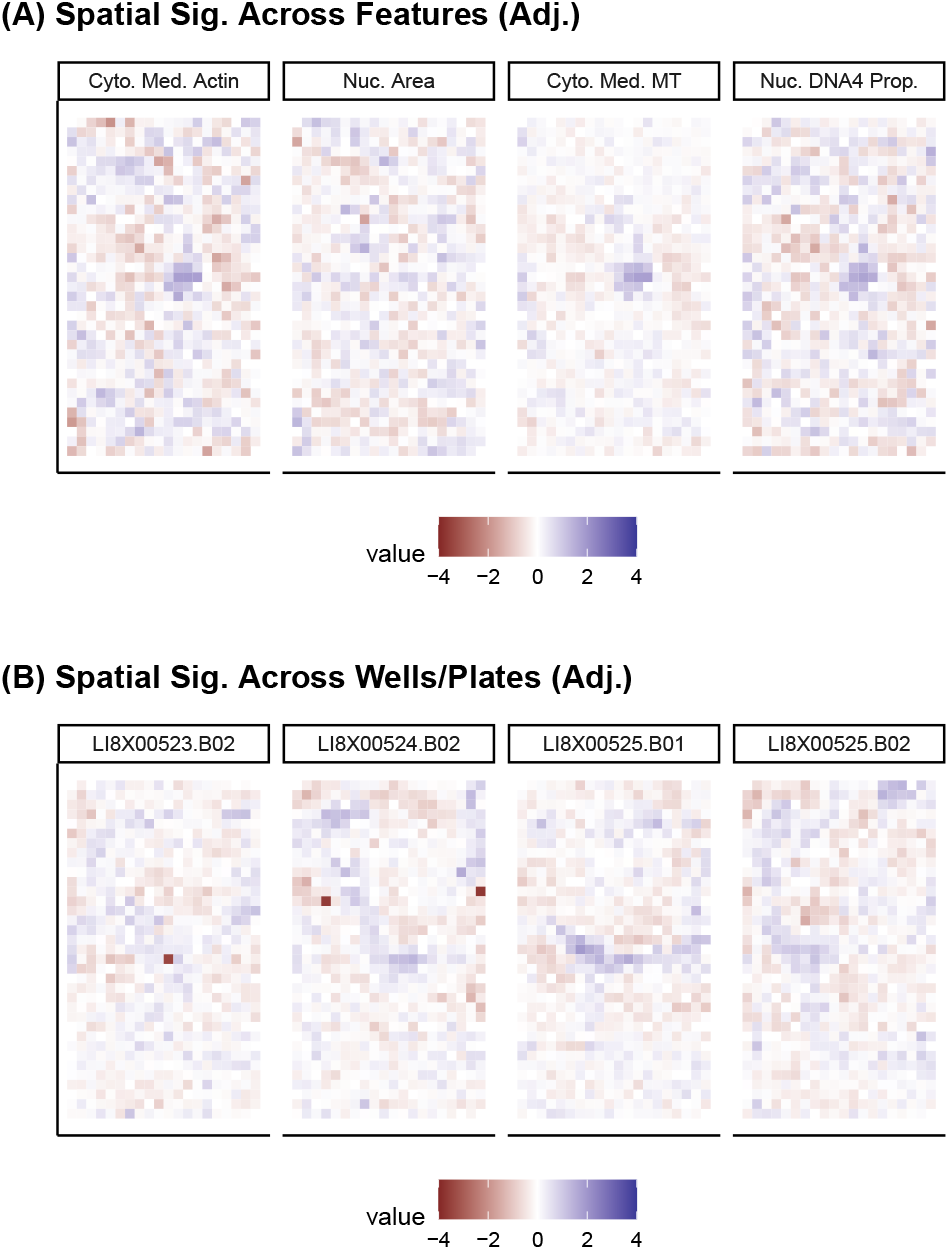
Similar to Figure 2 but after removing unwanted variation using our procedure.

### 3.0.3 Biological Signal is Preserved

Thrombospondin (THBS) is an ECMp that has been shown to have a particularly strong effect on MCF10A cells (Hunt *et al*., 2020). Consequently, we consider how prominent the signal of THBS is in our data before and after adjustment. To do this we display scatter plots in Figure 5 of the first two principal components before and after the normalization adjustment. This figure focuses on the nuclei DNA2N proportion feature as an illustrative example. (Figure 6 will more broadly consider recovery of the THBS in other features). In subplot (A) of this figure we color the points as either having been treated by THBS (in blue) or having been treated by some other ECMp (in red). While THBS stands out somewhat in the unadjusted plot, the clustering we see is weak and entirely contained as a second-order signal in the second principal component. Conversely, after adjustment we can see that not only does THBS prominently cluster away from the other ECMps but this clustering is also a first-order signal.

**Figure 5:**
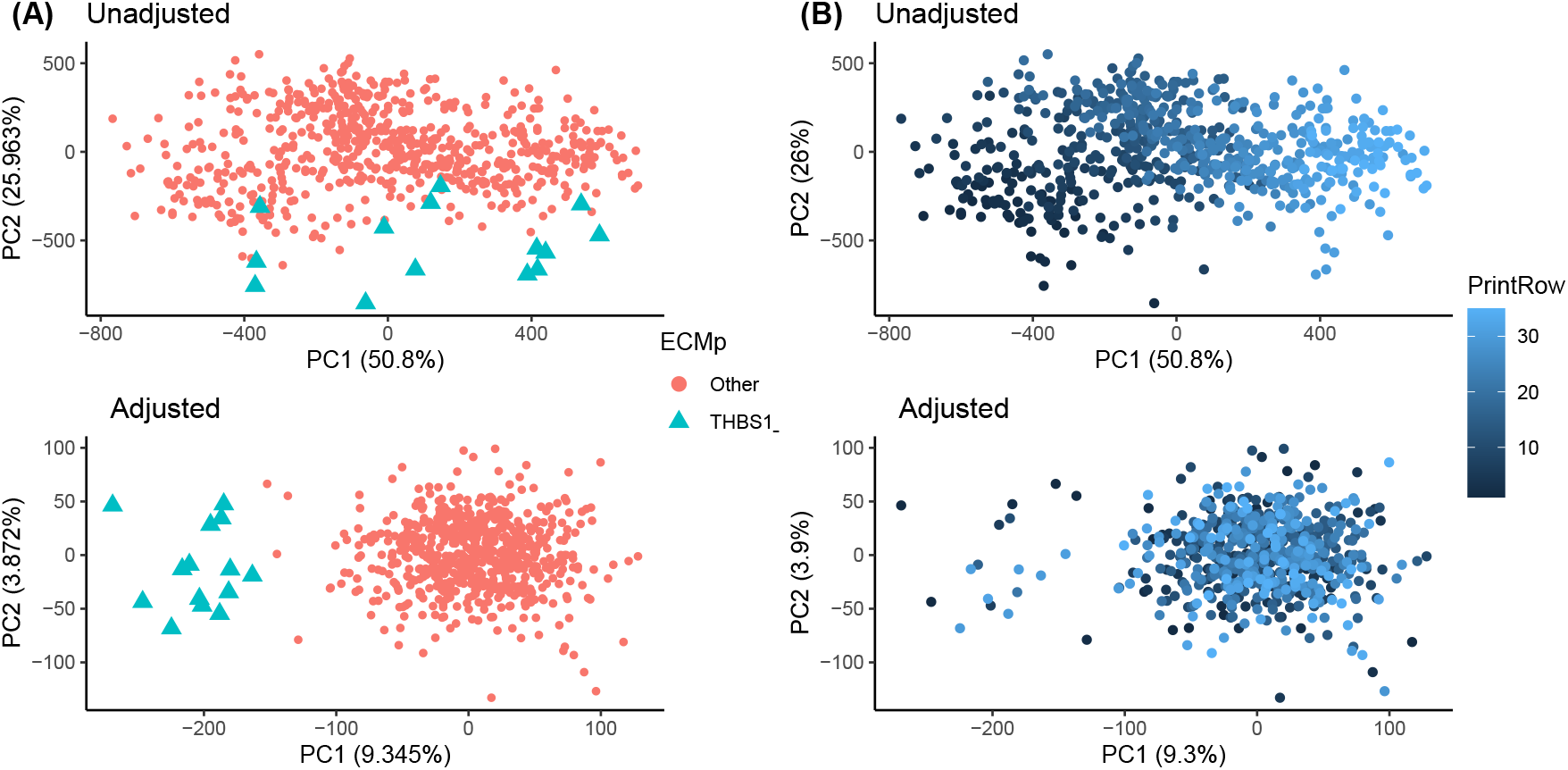
(A) Scatter plot of the first two PCs. Each point is a spot. The blue colored spots are those spots containing THBS. The top is before adjustment for unwanted variation and the bottom is after removing the unwanted effects. (B) similar to subplot (A) but the color indicates physical row of the well layout.

**Figure 6:**
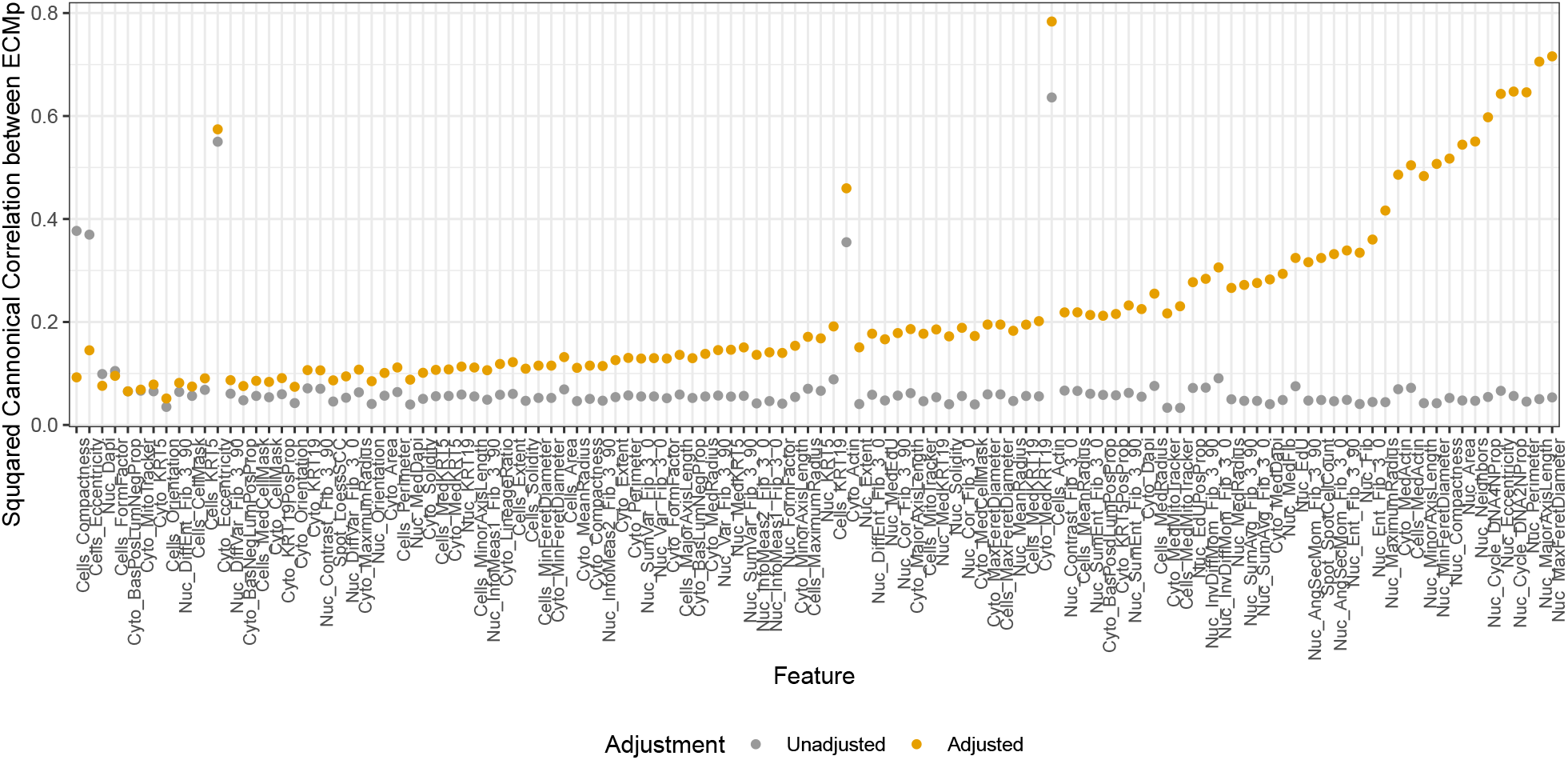
Squared canonical correlation between ECMp indicator variables and the first principal component of the feature matrix. This is calculated after adjustment (yellow) and before adjustment (gray). The ordering is by difference between pre- and post-adjustment canonical correlations. This figure illustrates that our adjustment allows the biological ECMp effect to be more prominently captured as a first-order effect in the data.

In Figure 5 (B) we plot the same points as in subplot (A) but color the points by the physical row location of the spot. We can see that before adjustment the first principal component largely captures a row-wise spatial effect. Conversely, after adjustment this row effect disappears. This illustrates the action of our approach which identifies the spatial effect and removes it. Consequently, as seen in the bottom of Fig 5 (A), we see the biological signal of THBS coming to the forefront and no longer see a color gradient in subplot (B). While in cases like this one may be tempted to simply remove the first principal component from analysis, this will not be a generalizable approach as the first principal component may not always be unwanted. Rather, our approach leverages replication to identify the truly unwanted effects.

In Figure 6 we summarize the results for all features comparing no adjustment to removing the unwanted effects using our approach. Here, we plot the squared canonical correlation between the 45 ECMp indicator covariates and the first principal component. This captures how much the signal due to the ECMPs is captured by the first PC and thus provides a measure of how prominent the desired ECMp signal is in the data before and after adjustment. From this figure we can see that without removing the unwanted effects (in gray) the first PC does not capture the ECMp signal in the data. This is likely because a dominating effect present in the data is spatial like that of Figure 5 (B). Conversely, after identifying and removing the unwanted effects we find that the biological ECMp signal more strongly shows up as a first-order signal since it is captured more by this first principal component (as measured by the canonical correlation). This figure shows that in nearly every case the adjustment we propose helps highlight the biology and minimize the unwanted artifacts. Our approach to normalizing MEMAs thus removes the unwanted spatial effects and thereby enhances the remaining biological signal.

## 4 Conclusion

High-throughput imaged-based profiling of cells and tissues is an increasingly widely used technique. Like all biological platforms, imaging assays are not immune to unwanted technical effects. In this work, we have focused on MEMAs to illustrate a design-based approach to identify and remove such unwanted effects. Our approach leverages replication at several levels of the experimental design to separate the desired biological signal from the unwanted effects.

While we demonstrate the efficacy of the proposed approach in the context of a specific MEMA experiment, the method is very general. Our approach will work for any set of feature matrices *Y* ^*i*^ with any design having replication across both the rows and the columns. The row replication identifies the unwanted effects *S* while the column replicates help find the weights *W*^*i*^ that govern how much of the unwanted effects to remove. Implicitly, our approach requires that the unwanted effects vary across these replicates. Otherwise, we will not be able to identify and remove them.

While for MEMAs the rows in our design correspond to wells and the columns correspond to spots, for other image-based experiments the rows and columns may correspond to different facets of the design that are replicated. Thus, while our application on MEMA data shows that the approach removes unwanted spatial effects, the generality of the approach we propose suggests it is applicable to removing many other types of unwanted effects from other platforms. Indeed, the approach requires little prior characterization of the unwanted effects except that it can be captured in a low-dimensional space.

While it is important for our approach that the entire replication structure is observed for some subset of the feature matrices, it is not required that this structure is completely observed for all features. As long as we have some features without rows systematically missing we can learn the unwanted effects from these features and subsequently remove them from all features. In our case this is done with the DAPI-derived features.

We believe the approach we describe may be broadly applicable to many other recently developed imaging techniques, including multiplexed and cyclic immunofluorescence and Cell Painting, and to other other multi-well plate experiments like 96- or 384-well plates. Like the MEMA data we present in this paper, it is often straight-forward to build replication into the design for these experiments, e.g. through a repeated cycle of staining, imaging, and dye inactivation or through a randomization of treatments across the rows and columns of the plates. Consequently these types of data are likely highly amenable to similar replication-based normalization. More broadly, the approach we advocate helps demonstrate the utility of replication in experimental design and highlights how careful use replication can substantially improve data quality and subsequently biological interpretation.

## 5 Data Availability and Software

Raw data is available on synapse (syn2862345), analysis code is on github (gjhunt/mema norm), a Docker image is available on dockerhub (gjhunt/memanorm).

## References

Ahmed Raza, S. E. et al. (2016). Robust normalization protocols for multiplexed fluorescence bioimage analysis. BioData Mining, 9(1), 11.

Akturk, G. et al. (2020). Multiplexed Immunohistochemical Consecutive Staining on Single Slide (MIC-SSS): Multiplexed Chromogenic IHC Assay for High-Dimensional Tissue Analysis. Methods in molecular biology (Clifton, N.J.), 2055, 497–519.

Bankhead, P. et al. (2017). QuPath: Open source software for digital pathology image analysis. Scientific Reports 2017 7:1, 7(1), 1–7.

Bray, M.-A. et al. (2016). Cell Painting, a high-content image-based assay for morphological profiling using multiplexed fluorescent dyes. Nature Protocols 2016 11:9, 11(9), 1757–1774.

Carpenter, A. E. et al. (2006). CellProfiler: image analysis software for identifying and quantifying cell phenotypes. Genome Biology 2006 7:10, 7(10), 1–11.

Chang, Y. H. et al. (2020). RESTORE: Robust intEnSiTy nORmalization mEthod for multiplexed imaging. Communications Biology, 3(1), 111.

Collins, T. J. (2007). Imagej for microscopy. BioTechniques, 43(1S), S25–S30. PMID: 17936939.

Drew, C. P. and Shieh, W.-J. (2015). Chapter ten - immunohistochemistry. In C. Rupprecht and T. Nagarajan, editors, Current Laboratory Techniques in Rabies Diagnosis, Research and Prevention, Volume 2, pages 109–115. Academic Press.

Frisch, R. and Waugh, F. V. (1933). Partial Time Regressions as Compared with Individual Trends. Econometrica, 1(4), 387–401.

Hamilton, N. (2009). Quantification and its applications in fluorescent microscopy imaging. Traffic, 10(8), 951–961.

Harris, C. et al. (2021). Quantifying and correcting slide-to-slide variation in multiplexed immunofluorescence images. bioRxiv.

Heilemann, M. (2012). 2.4 super-resolution microscopy. In E. H. Egelman, editor, Comprehensive Biophysics, pages 39–58. Elsevier, Amsterdam.

Hunt, G. J. et al. (2020). Automatic transformation and integration to improve visualization and discovery of latent effects in imaging data. Journal of Computational and Graphical Statistics, 29(4), 929–941.

Huwyler, J. and Pardridge, W. M. (1998). Examination of blood-brain barrier transferrin receptor by confocal fluorescent microscopy of unfixed isolated rat brain capillaries. Journal of Neurochemistry, 70(2), 883–886.

Ishizawa, T. et al. (2009). Real-time identification of liver cancers by using indocyanine green fluorescent imaging. Cancer, 115(11), 2491–2504.

Lin, C.-H. et al. (2012). Fabrication and Use of MicroEnvironment microArrays (MEArrays). Journal of Visualized Experiments, (68), 1–7.

Lin, J.-R. et al. (2016). Cyclic immunofluorescence (cycif), a highly multiplexed method for single-cell imaging. Current Protocols in Chemical Biology, 8(4), 251–264.

Smith, R. et al. (2019). Using Microarrays to Interrogate Microenvironmental Impact on Cellular Phenotypes in Cancer. Journal of visualized experiments : JoVE.

Sommer, C. et al. (2011). Ilastik: Interactive learning and segmentation toolkit. In 2011 IEEE International Symposium on Biomedical Imaging: From Nano to Macro, pages 230–233.

Sood, A. et al. (2020). Comparison of Multiplexed Immunofluorescence Imaging to Chromogenic Immunohistochemistry of Skin Biomarkers in Response to Monkeypox Virus Infection. Viruses, 12(8).

Tan, W. C. C. et al. (2020). Overview of multiplex immunohistochemistry/immunofluorescence techniques in the era of cancer immunotherapy. Cancer Communications, 40(4), 135.

Watson, S. S. et al. (2018). Microenvironment-Mediated Mechanisms of Resistance to HER2 Inhibitors Differ between HER2+ Breast Cancer Subtypes. Cell Systems, 6, 329––342.e6.

